# Maternal brain gain: enlarged representation of the peripersonal space in pregnancy

**DOI:** 10.1101/492017

**Authors:** Flavia Cardini, Natalie Fatemi-Ghomi, Katarzyna Gajewska-Knapik, Victoria Gooch, Jane Elizabeth Aspell

**Affiliations:** School of Psychology and Sport Science, Anglia Ruskin University, East Road, CB1 1PT, Cambridge, UK.; Department of Obstetrics and Gynaecology, Cambridge University Hospitals NHS FT, Cambridge Biomedical Campus, CB2 0QQ, Cambridge, UK.

## Abstract

Our ability to maintain a coherent bodily self despite continuous changes within and outside our body relies on the highly flexible multisensory representation of the body, and of the space surrounding it: the peripersonal space (PPS).

The aim of our study was to investigate whether during pregnancy - when extremely rapid changes in body size and shape occur - a likewise rapid plastic reorganization of the neural representation of the PPS occurs. We used an audio-tactile integration task to measure the PPS boundary at different stages of pregnancy. We found that in the second trimester of pregnancy and postpartum women did not show differences in their PPS size as compared to the control group (non-pregnant women). However, in the third trimester the PPS was larger than the controls’ PPS and the shift between representation of near and far space was more gradual. We therefore conclude that during pregnancy the brain adapts to the sudden bodily changes, by expanding the representation of the space around the body. This may represent a mechanism to protect the vulnerable abdomen from injury from surrounding objects.

## Introduction

Pregnancy is an exceptional and temporary condition in a woman’s life, when rapid changes occur in the body - both internally, and externally. During this short period a woman’s body rapidly changes in size and shape while new and compelling signals arise from inside. During pregnancy, the maternal brain’s representation of the internal body would be expected to change, firstly because of the entirely new signals due to foetal growth and movements, and the consequent abdominal changes, and secondly because the maternal brain needs to monitor a new entity: the foetal body. Additionally, the maternal brain should update the representation of the external body to accommodate its new dimensions, for example to ensure that the mother does not sustain injury to her protruding, vulnerable abdomen from nearby objects. Given the magnitude and rapidity of the bodily changes occurring during pregnancy, one would expect them to be coupled with a likewise rapid plastic reorganization of the neural representation of both the body and the space surrounding it.

Neuropsychological, neuroimaging and behavioural data have highlighted the presence of both slow, long-term neuroplastic changes that are associated with expertise^1^, or that simply reflect developmental processes^2^, and more rapid updates of the body representation that occur during the continuous interaction with the external world^3,4^. For example, blind subjects who use a cane to navigate, respond to auditory stimuli arising from locations near the tip of the cane in a similar way to those presented near their hand. This is likely due to an expansion of visuo-tactile neuronal receptive fields around the hand to represent the space surrounding the cane, suggesting that when we become expert at using a tool, we perceive it as part of our own body^5,6^.

The space around our body - the so-called peripersonal space (PPS)^7,8^ - is the interface between the body and the environment since it is the area of space where physical interactions with the external world take place. This special region of the space is constantly monitored by the brain because only within its boundaries can we reach and act upon objects. Recently, it has been shown that the representation of the PPS is not stable, but rather flexible^9,10^. For example, being in proximity to an individual we have previously co-operated with induces an expansion of our PPS towards that person, so that it grows to encompass the space between ourselves and the other^11^. More recent results show that sharing a sensory experience with another person induces a remapping of the other’s PPS onto one’s own PPS^12^. Finally, experimental evidence has demonstrated an expansion of the PPS representation after tool use: when an individual acts upon far space with a tool, their brain’s representation of near and far space changes, with the far space being treated as near space^13–15^. This effect was first described in monkeys^16^ and later, a similar mechanism was observed also in humans^17^. In brief, after training with a tool, plastic changes both to the representation of the dimensions of the body part acting upon the tool - and of the PPS, were observed. Taken together, these studies demonstrate how plastic the representation of the space surrounding one’s body can be.

Since there are no reports to date (according to a recent review paper^18^), about the brain changes in own-body representation that very likely occur during pregnancy this study investigated the changes in the representation of the space around the body in pregnant women. In order to test whether the boundary between near and far space is modulated in pregnancy we measured the size of the PPS using an audio-tactile Reaction Time (RT) task^19^ in pregnant women at three stages: at an early stage of their gestational period (~20^th^ week of gestational period), at a later stage (~34^th^ week) and a few weeks (~8 weeks) postpartum and we compared each measure with that taken from a control group of non-pregnant women. We expected to find that the representation of the PPS would expand with advancing gestation and that it would shrink back to its former size after birth while it would remain unchanged over time for non-pregnant women.

## Results

Participants’ RTs to tactile stimulation of the abdomen were recorded for each trial at each tapping delay after sound onset. Trials that were faster or slower than 2.5 SD of their average RT for that onset delay were removed (< 5% of total trials). To investigate whether PPS representation changes in pregnancy, mean RTs to the tactile stimulus administered at the different delays were calculated and compared between the two groups in each testing session.

For each testing session a 5×2 mixed ANOVA was run with Delay (D1, D2, D3, D4, D5 with D1 = farthest distance from participant and D5 = closest distance) as the within-subjects factor and Group (pregnant women vs non-pregnant controls) as the between-subjects factor.

For testing session 1 (i.e. when pregnant participants were at the ~20^th^ week of their gestational period), a main effect of Delay (*F*_(4,216)_ = 47.16, *p* < .001, *ηp*^2^ = .466) was found. Main effects of Group and Delay x Group interaction were non-significant. To analyse the significant main effect of Delay further, we ran four two-tailed paired samples t-tests to compare RTs between each consecutive delay. The alpha-level was Bonferroni corrected for multiple comparisons to *p* = .012. Post-hoc tests revealed non-significant difference between RTs at D4 (M = 1050.87, SE = 11.76) and D5 (M = 1042.88, SE = 10.76), (*t*_(55)_ = 1.82, *p* = .074), whereas RTs significantly differ between D3 (M = 1069.61, SE = 13.02) and D4 (*t*_(55)_ = 3.92, *p* < .001), between D2 (M = 1141.85, SE = 13.31) and D3 (*t*_(55)_ = 5.86, *p* < .001) and between D1 (M = 1182.44, SE = 16.01) and D2 (*t*_(55)_ = 4.19, *p* < .001). See Figure 1 (graph “a”).

For testing session 2 (i.e. when pregnant women were at the ~34^th^ week of their gestational period), a significant main effect of Delay (*F*_(4,172)_ = 63.69, *p*< .001, *ηp*^2^ = .597) and a significant Group x Delay interaction (*F* _(4,172)_ = 4.62; *p* = .012, *ηp*^2^ = .097) were found. No main effect of Group was found. Bonferroni-corrected (*p* = .012) two-tailed paired-sample t-tests showed a non significant difference between RTs at D4 (M = 1034.93, SE = 9.70) and D5 (M = 1019.42, SE = 8.34), (*t*_(44)_ = 2.58, *p* = .013) and non significant difference between RTs at D3 (M = 1054.46, SE = 9.89) and D4 (*t*_(44)_ = 2.43, *p* = .019), whereas a significant difference was found between RTs at D2 (M = 1101.43, SE = 10.49) and D3 (*t*_(44)_ = 8.78, *p* < .001), and between RTs at D1 (M = 1146.25, SE = 14.27) and D2 (*t*_(44)_ = 4.58, *p* < .001). Additionally, to study the source of the significant two-way interaction, we ran four post-hoc Holm-Bonferroni corrected paired samples t-tests for each group. The aim of these analyses was to identify the critical distance at which the sound speeded up tactile RTs (that can be taken as a proxy of the boundary of PPS, see^11,19^), and to test whether this distance differed between the two groups. In the Control group a non-significant difference between RTs at D4 (M = 1017.11, SE = 16.71) and D5 (M = 998.71, SE = 13.03) was found, (*t*_(16)_ = 1.87, *p* = .079), whereas RTs at D3 (M = 1061.39, SE = 17.86) were slower than at D4 (*t*_(16)_ = 2.91, *p* = .010), RTs at D2 (M = 1100.99, SE = 20.28) were slower than RTs at D3 (*t*_(16)_ = 4.46, *p* < .001) and RTs at D1 (M = 1170.56, SE = 30.50) were slower than RTs at D2 (*t*_(16)_ = 3.78, *p* = .002).

Interestingly, in the pregnant group, t-tests showed a non-significant difference between RTs at D4 (M = 1045.76, SE = 11.59) and D5 (M = 1031.99, SE = 10.27), (*t*_(27)_ = 1.78, *p* = .085) and between RTs at D3 (M = 1050.25, SE = 11.81) and D4,(*t*_(27)_ = .56, *p* = .576), whereas RTs at D2 (M = 1101.70, SE = 11.86) were slower than RTs at D3 (*t*_(27)_ = 7.69, *p* < .001), and RTs at D1 (M = 1131.50, SE = 13.32) were slower than RTs at D2 (*t*_(27)_ = 2.88, *p* = .008). See Figure 1 (graph “b”).

This means that in the two groups, tactile processing is differently boosted by co-occurring sounds, with a facilitation effect of sound on RTs occurring between D2 and D3 for pregnant women, and between D3 and D4 for the control group. By taking the critical distance at which the sound speeds up tactile RTs as a proxy of the PPS boundary, we can conclude that the PPS size of pregnant women at a late stage of gestational period is larger than that in non-pregnant women.

For testing session 3 (i.e. ~8 weeks postpartum), a main effect of Delay (*F* _(4,132)_ = 39.87; p < .001, *ηp*^2^ = .547) was found. Main effects of Group and the Delay x Group interaction were not significant. In order to follow-up the significant main effect of Delay, Bonferroni-corrected post-hoc t-tests revealed non significant difference between RTs at D4 (M = 1013.71, SE = 9.43) and D5 (M = 1004.28, SE = 9.34), (*t*_(34)_ = 1.62, *p* = .113), whereas RTs significantly differ between D3 (M = 1036.16, SE = 8.37) and D4 (*t*_(34)_ = 2.99, *p* = .005), between D2 (M = 1070.02, SE = 10.99) and D3 (*t*_(34)_ = 3.28, *p* = .002) and between D1 (M = 1122.54, SE = 14.56) and D2 (*t*_(34)_ = 4.99, *p* < .001). See Figure 1 (graph “c”).

Additionally, we fitted the data of the two groups to a linear function to assess any group differences between the steepness of the slopes of the PPS gradient. The linear function was described by the following equation:

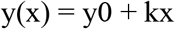

where x represents the delay, y the RT, y0 the intercept at x = 0, and k the slope of the function. Due to bad fitting (R2 < .20), two participants in testing session 1, three participants in testing session 2 and three participants in testing session 3 were excluded from further analysis. For each testing session, we then compared the slope of the function between the two groups. In testing session 1, no significant difference was observed between the slope of the linear function computed in the pregnant women group and that computed in the control group, (*t*_(52)_ = .316, *p* = .75).

Importantly, in testing session 2 the slopes significantly differed between the two groups, with the slope for the pregnant women less steep (k = -.046) than that in the control group (k = -.07), (*t*_(40)_ = 2.15, *p* = .03). In testing session 3, no difference was found between the slopes of the two groups (*t*_(30)_ = 1.07, *p* = .29). See Figure 1.

**Figure 1.**
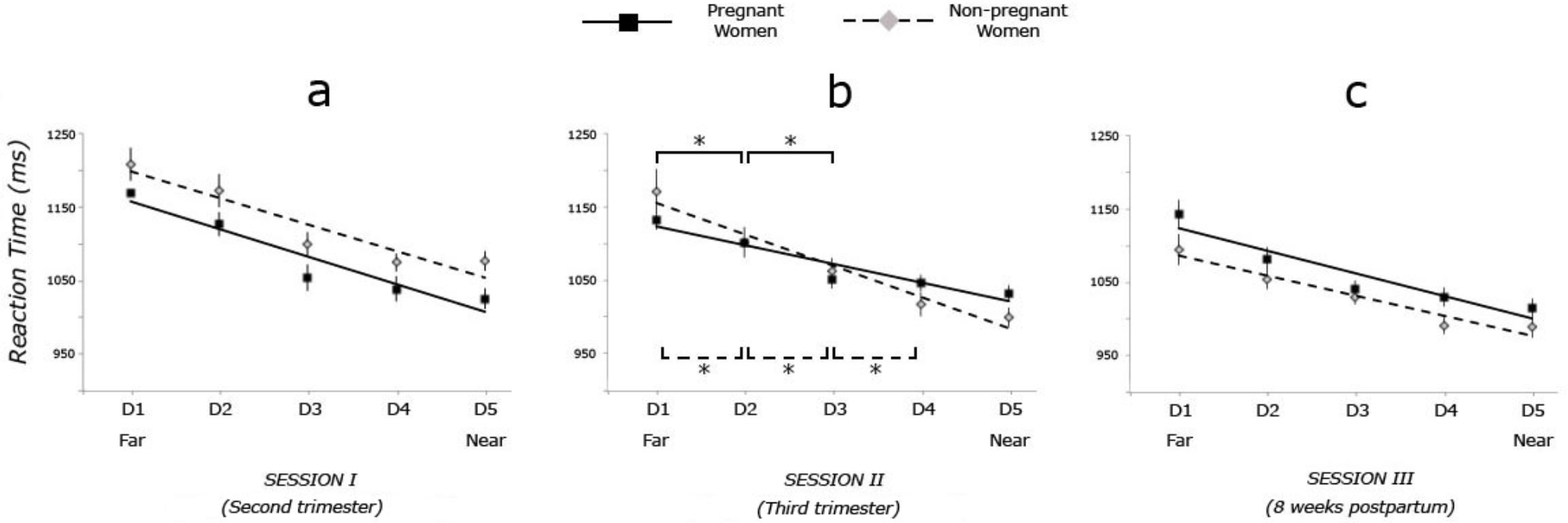
Results of the audio-tactile task. Plots of performance of two groups on the audio-tactile PPS task, when pregnant women were in their second trimester (i.e. at their ~20^th^ week of gestation period, graph “a”), in their third trimester (i.e. at their ~34^th^ week, graph “b”) and ~8 weeks postpartum (graph “c”). Mean reaction times (RTs) to tactile stimuli (in ms, y axis) were measured at five distinct time periods, during which an auditory stimulus was perceived moving towards the participant’s own body. Error bars reflect standard error of the mean, and asterisk indicates p-value < .05, two-tailed. Solid lines refer to pregnant women’s performance and dashed lines to that of non-pregnant women.

## Discussion

The aim of the present study was to investigate whether changes in the representation of peripersonal space occur during pregnancy. In line with previous research^11,19^, in this study we took the critical distance at which coincident sounds speed up tactile RTs as a proxy of the PPS boundary, and we used the slope of the PPS gradient as an index of the speed in the transition from peri- to extrapersonal space. Our results show that at a late stage of pregnancy (i.e. at around the 34^th^ week of gestational period) significant changes occur in the representation of the space around the body. In particular, both the size of the PPS increases and the gradient in the transition between near and far space becomes shallower. This PPS reshaping is not observed at an earlier stage of pregnancy nor a few weeks postpartum, when the size and shape of the PPS is comparable to that of non-pregnant women. These results therefore suggest that only when the body undergoes significantly large changes, does the brain adapt the representation of the surrounding space accordingly. Our findings are in line with several studies revealing how rapidly the representation of the PPS adapts to experimentally-induced changes. For example, in a recent study Canzoneri and colleagues investigated the effect of tool use on both body and PPS representations^17^. After twenty minutes of using a long tool to reach far objects, participants perceived the shape of their forearm as becoming similar to the one of the tool, i.e. narrower and longer, as compared to before the tool-use. At the same time, participants’ representation of the PPS expanded towards the tip of the tool, as to incorporate it into one’s own body representation.

In the cognitive neuroscience literature two main, non-mutually exclusive functional roles of the PPS have been identified so far: the PPS has been described as the sensorimotor interface for goal-oriented actions^7,20^; it also acts as a “protective bubble” that keeps a margin of safety around the body, with the aim of coordinating defensive behaviours against potentially dangerous stimuli^21^. Several studies have confirmed this defensive role of the PPS. For example, Taffau and Viaud-Delmon showed that in response to an approaching sound of a barking dog, participants’ PPS increased in size. This effect was specifically observed in cynophobic people - i.e. people with phobia for dogs – therefore leading to the conclusion that the PPS expands only when the need of protecting the body from an approaching potentially threatening stimulus emerges^22^. Similarly, in a time-to-collision study, Vagnoni and colleagues showed that the looming image of a feared animal - such as a spider or snake - is perceived as colliding with the observer’s body sooner than a neutral looming image, indicating an expansion of the observer’s PPS boundaries only in the presence of a threatening stimulus^23^. The present results seem in line with the defensive account of the PPS. Pregnancy involves massive and rapid changes in the body both externally – as the body suddenly assumes new dimensions – and internally – while the foetus is growing. As a consequence, the maternal brain has to adequately react to such critical changes. Therefore, we suggest that the observed expansion of the PPS at a late stage of pregnancy might be aimed at protecting the vulnerable abdomen – and the new entity held within it – during the mother’s daily interaction with the external environment.

Importantly, our study not only shows an increased size of the PPS in late pregnancy, but also a shallower gradient with which the perceived space transitions from peripersonal to extrapersonal. In a recent review paper, Noel and colleagues suggest that the shallowness or steepness of the PPS reflect the gradient in the boundary between one’s own body and the others’^24^. Whereas a slow transition space, indexed by a shallow PPS representation, is related with a weak self-other distinction, a steep PPS representation seems in line with a sharp and inflexible self-other boundary. Although this theoretical account of the PPS is build up on evidence from studies on schizophrenia and Autism Spectrum Disorder^25,26^ (see also Mul et al., under review), the suggested neurocognitive mechanisms involved, could plausibly explain our results as well.

The shallower slope of the PPS gradient observed at the late stage of pregnancy seems therefore to indicate a weaker and more variable sense of body boundary, perhaps caused by the inability of the brain to accurately keep track of the fast body changes. Interestingly, the PPS has been defined as a “stochastic bubble” where computations about the probability of the body interacting with external objects continuously occur^27^. The wider margin of the “safety zone” around the body observed late in pregnancy could be a consequence of the brain’s reduced ability to accurately compute the exact spatial location of an external stimulus with respect to one’s own – rapidly growing - body. Therefore, given this enhanced uncertainty, the brain starts treating stimuli - usually perceived as far away – as if located in the near space. The current results are in line with previous qualitative studies investigating the experience of one’s own body during pregnancy^28,29,30^. By interviewing pregnant women at different stages of their gestational period, the authors identified an interesting theme: some women reported a sense of disrupted body boundaries and confusion in their bodies’ separation from both the fetus and the external world.

An alternative – although highly speculative – explanation for the shallower PPS gradient takes into consideration the role of empathy. It has been suggested that whereas a steep and inflexible boundary between self and others prevents social communication and the ability to adequately understand the others’ mental and physical states, a shallower gradient seems to facilitate the process of empathising with others^24^. Given this idea of a relationship between the steepness of the PPS gradient and empathic traits, we might expect that the boundary between a pregnant woman’s body and others’ will be expanded, with the aim of facilitating bonding with the future newborn. Indeed, maternal behaviour - critical for an infant’s survival^31,32^ – strongly depends on the mother’s ability to promptly understand the infant’s cues, predict their needs and adequately react to them^33^, i.e. her empathic ability. However, this explanation has a highly speculative nature, as - according to a recent review^34^ on empathy in pregnancy and in the postpartum period - no clear evidence yet exists to support our hypothesis.

To conclude, research on the neural representation of the body usually relies on the generation of transient illusory effects, such as experimentally-induced changes in the perception of one’s own body and its surrounding space (see the Rubber Hand Illusion^3^, the Full Body Illusion^35^). Additional evidence on the mechanisms underlying the representation of one’s own body and the PPS is provided by investigations of the slower and more long-lasting plastic changes in the body representation following training and learning (see effect of tool-use training on the PPS^9^). Although such experimentally-induced changes are needed to shed light on the different sources of information that contribute to the representation of the body, bodily illusions cannot reveal whether natural changes in body configuration are coupled with plastic changes in the cortical representation of the body. With this study, for the first time, we overcame this limitation, by investigating an exquisite case of non-experimentally induced change in body size and its effect on the mental representation of the body and its surrounding space. Rapid changes in the representation of PPS may also occur – but have yet to be studied – as a result of other developmental processes, e.g. during growth spurts. We predict that changes in PPS representation would also occur following large increases or decreases in abdomen size due to weight gain or loss. Pregnancy, however, might result in more rapid PPS changes than those arising from weight gain, because of the greater vulnerability of the foetus and the strong evolutionary imperative to protect it. Further investigation of brain plasticity induced by the bodily changes accompanying pregnancy is likely to be a fertile avenue for future research.

## Methods

### Participants

37 pregnant women (Age range = 21-43; M_age_ = 31; SD = 4,8) and 19 controls (Age range = 21-43; M_age_ = 31,4, SD =7,3) took part in the first testing session.

28 pregnant women (Age range = 23-43; M_age_ = 31,8; SD = 4,7) and 17 controls (Age range = 21-43; M_age_ = 30,5, SD =7,3) took part in the second testing session.

20 pregnant women (Age range = 23-43; M_age_ = 32,4; SD = 5,4) and 15 controls (Age range = 21-43; M_age_ = 30,6; SD = 6,9) took part in the third testing session. Due to a high dropout rate in the pregnant women’s group between the first and the second session, extra participants (N=4) were recruited late in their pregnancy, therefore they took part only in the second and third sessions.

Procedures were approved by the East of England - Cambridgeshire and Hertfordshire Research Ethics Committee (15/EE/0294) and were in accordance with the principles of the Declaration of Helsinki.

Participants were recruited by the midwife (NF-G) involved in the project at Addenbrooke’s Hospital in Cambridge (UK), provided written informed consent and were reimbursed for participation.

For the experimental group, only participants with low-risk singleton pregnancy and with BMI below 28 – as measured at the booking scan at 12 weeks - were allowed to take part in the study. No participants reported any neurological conditions.

### Procedure

In order to assess any changes in the boundaries of PPS during pregnancy, pregnant women in their second trimester - Mean week of gestational period = 20^th^ (SD=3.5) - in their third trimester - Mean week of gestational period = 34^th^ (SD=0.9) - and approximately 8 weeks postpartum (SD=1.5) were tested. Additionally, a control group of non-pregnant women was tested in each testing session. Participants were asked to perform an audio-tactile task adapted from Canzoneri et al.^19^, where participants sat blindfolded with their left arm resting on a response box on a table beside them. On each trial, a task-irrelevant sound was presented for 3000ms. The sound was generated by two loudspeakers: one was placed on the table close to the participant’s hand and the other one, 1m further away. Auditory stimuli were samples of pink-noise, at 44.1 kHz. Sound intensity was manipulated using Audacity software, so that the sound had an exponentially rising acoustic intensity from 55 to 70 dB Sound Pressure Level (SPL) as measured with an audiometer positioned at the participant’s ear at the beginning of the experiment. The sound was a combination of two identical samples of pink noise, one of increasing and the other one of decreasing intensity, emitted by the near and far loudspeakers respectively. Both loudspeakers were activated simultaneously, but whereas the far loudspeaker activated at the maximum intensity and then its intensity decreased up to silence along the trial, the near loudspeaker activated at the minimum intensity, and then its intensity increased up to the maximum value along the trial. In this way, participants had the impression of a sound source moving from the far to the near loudspeaker, i.e. towards their own body.

While the sound was played, the participant’s abdomen was stimulated by using a custom built small tapper attached to it. In each trial, the tactile stimulation could be delivered at any of five possible delays from the onset of the sound: D1, tactile stimulation administered 300ms after the sound onset; D2, tactile stimulation administered 800ms after the sound onset; D3, tactile stimulation administered 1500ms after the sound onset; D4, tactile stimulation administered 2200ms after the sound onset; D5, tactile stimulation administered 2500ms after the sound onset. In this way, tactile stimulation occurred when the sound source was perceived at different locations with respect to the body: i.e., far from the participant’s body - at short temporal delays - and gradually closer to the participant’s body, as the temporal delays increased (see Figure 2). Arduino board and LabView 8 (National Instruments, Austin, TX) software were used to control the sound, onset delay of the tap and record reaction times (RTs). Participants were asked to respond as quickly as possible to the tactile stimulation by pressing a key with their left hand and to ignore the sound. Ten trials for each temporal delay and 15 catch trials (where the sound was played but not tactile stimulation delivered) were presented in a random order, resulting in a total of 65 trials. The task lasted approximately 7 minutes. As sounds facilitate tactile RTs only when presented close to the body^1^, we expected RTs to progressively decrease as the sound was approaching. The critical distance where the sound speeds up tactile RTs can be taken as a proxy of the PPS boundary.

**Figure 2.**
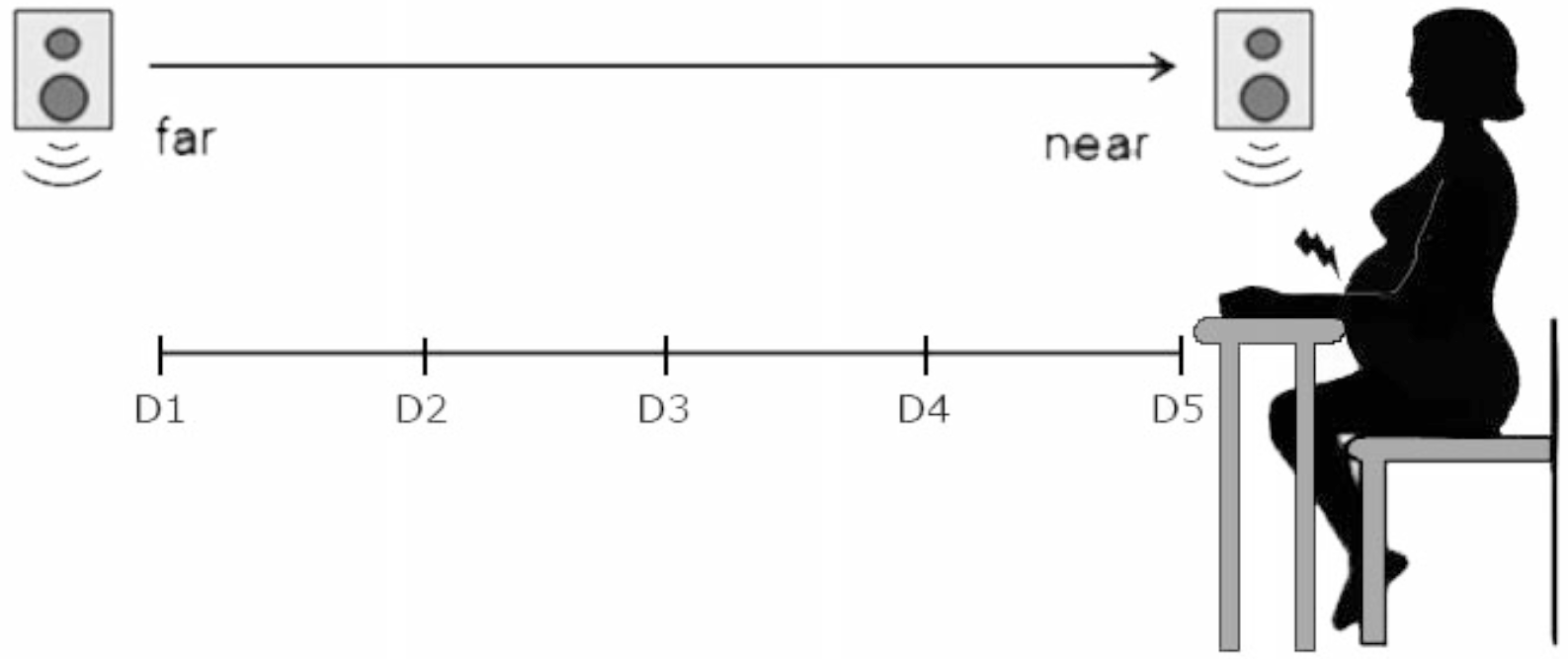
Experimental set-up of the audio-tactile task. Participants made speeded button-press responses to tactile stimuli (mechanical stimulation from a tapper attached to the abdomen), whilst seated blindfolded. During each trial, a 3sec sound was played via two loudspeakers, which gave the perception of a sound travelling towards the participant’s body. The tactile stimuli could be presented at one of five time-points during the sound, which corresponded to five perceived distances from the participant’s body ranging from far (D1 = 300ms) to near (D5 = 2500ms) the participant. RTs to the tactile stimulus were recorded.

## Acknowledgments

FC, JA and NF-G were supported by a Bial Foundation (Grant number 121/14) awarded to FC and JA.

## Author Contributions

FC and JA developed the study concept and the study design. FC, JA, VG and NF-G contributed to data collection. FC performed the data analysis. FC and JA contributed to the results interpretation. FC drafted the manuscript, and JA provided critical revisions. NF-G, KG-K and VG revised the manuscript. All authors approved the final version of the paper for submission.

## References

1. Serino, A., Bassolino, M., Farnè, A. & Làdavas, E. Extended multisensory space in blind cane users. Psychol. Sci. 18, 642–648 (2007).

2. Butler, D. L., Mattingley, J. B., Cunnington, R. & Suddendorf, T. Different neural processes accompany self-recognition in photographs across the lifespan: an ERP study using dizygotic twins. PLoS One 8, e72586 (2013).

3. Botvinick, M. & Cohen, J. Rubber hands ‘feel’ touch that eyes see. Nature 391, 756 (1998).

4. Longo, M. R. & Serino, A. Tool use induces complex and flexible plasticity of human body representations. Behav. Brain Sci. 35, 229–230 (2012).

5. Iriki, A., Tanaka, M., Obayashi, S. & Iwamura, Y. Self-images in the video monitor coded by monkey intraparietal neurons. Neurosci. Res. 40, 163–173 (2001).

6. Sengül, A. et al. Extending the body to virtual tools using a robotic surgical interface: evidence from the crossmodal congruency task. PLoS One 7, e49473 (2012).

7. Rizzolatti, G., Fadiga, L., Fogassi, L. & Gallese, V. The space around us. Science 277, 190–191 (1997).

8. di Pellegrino, G., Làdavas, E. & Farné, A. Seeing where your hands are. Nature 388, 730–730 (1997).

9. Cléry, J., Guipponi, O., Wardak, C. & Ben Hamed, S. Neuronal bases of peripersonal and extrapersonal spaces, their plasticity and their dynamics: knowns and unknowns. Neuropsychologia 70, 313–326 (2015).

10. Canzoneri, E., Marzolla, M., Amoresano, A., Verni, G. & Serino, A. Amputation and prosthesis implantation shape body and peripersonal space representations. Sci. Rep. 3, 2844 (2013).

11. Teneggi, C., Canzoneri, E., di Pellegrino, G. & Serino, A. Social modulation of peripersonal space boundaries. Curr. Biol. 23, 406–411 (2013).

12. Maister, L., Cardini, F., Zamariola, G., Serino, A. & Tsakiris, M. Your place or mine: Shared sensory experiences elicit a remapping of peripersonal space. Neuropsychologia 70, (2015).

13. Longo, M. R. & Lourenco, S. F. Space perception and body morphology: extent of near space scales with arm length. Exp. brain Res. 177, 285–290 (2007).

14. Patané, I., Iachini, T., Farnè, A. & Frassinetti, F. Disentangling Action from Social Space: Tool-Use Differently Shapes the Space around Us. PLoS One 11, e0154247 (2016).

15. Berti, A. & Frassinetti, F. When far becomes near: remapping of space by tool use. J. Cogn. Neurosci. 12, 415–420 (2000).

16. Iriki, A., Tanaka, M. & Iwamura, Y. Coding of modified body schema during tool use by macaque postcentral neurones. Neuroreport 7, 2325–30 (1996).

17. Canzoneri, E. et al. Tool-use reshapes the boundaries of body and peripersonal space representations. Exp. Brain Res. 228, 25–42 (2013).

18. Di Noto, P. M., Newman, L., Wall, S. & Einstein, G. The Hermunculus: What Is Known about the Representation of the Female Body in the Brain? Cereb. Cortex 23, 1005–1013 (2013).

19. Canzoneri, E., Magosso, E. & Serino, A. Dynamic Sounds Capture the Boundaries of Peripersonal Space Representation in Humans. PLoS One 7, e44306 (2012).

20. Brozzoli, C., Ehrsson, H. H. & Farnè, A. Multisensory representation of the space near the hand: from perception to action and interindividual interactions. Neuroscientist 20, 122–135 (2014).

21. Graziano, M. S. A. & Cooke, D. F. Parieto-frontal interactions, personal space, and defensive behavior. Neuropsychologia 44, 2621–2635 (2006).

22. Taffou, M. & Viaud-Delmon, I. Cynophobic Fear Adaptively Extends Peri-Personal Space. Front. Psychiatry 5, 122 (2014).

23. Vagnoni, E., Lourenco, S. F. & Longo, M. R. Threat modulates perception of looming visual stimuli. Curr. Biol. 22, R826–R827 (2012).

24. Noel, J.-P., Cascio, C. J., Wallace, M. T. & Park, S. The spatial self in schizophrenia and autism spectrum disorder. Schizophr. Res. 179, 8–12 (2017).

25. Kanner, L. Autistic disturbances of affective contact. Acta Paedopsychiatr. 35,100–136 (1968).

26. Nasrallah, H. A. Impaired mental proprioception in schizophrenia. Curr. Psychiatr. 11, 4–6 (2012).

27. Noel, J.-P., Blanke, O. & Serino, A. From multisensory integration in peripersonal space to bodily self-consciousness: from statistical regularities to statistical inference. Ann. N. Y. Acad. Sci. 1426, 146–165 (2018).

28. Hodgkinson, E. L., Smith, D. M. & Wittkowski, A. Women’s experiences of their pregnancy and postpartum body image: a systematic review and meta-synthesis. BMC Pregnancy Childbirth 14, 330 (2014).

29. Raphael-Leff, J. Pregnancy: the inside story. (Karnac Books, 2001).

30. Talmon, A. & Ginzburg, K. ‘Who does this body belong to?’ The development and psychometric evaluation of the Body Experience during Pregnancy Scale. Body Image 26, 19–28 (2018).

31. Feldman, R., Weller, A., Zagoory-Sharon, O. & Levine, A. Evidence for a neuroendocrinological foundation of human affiliation: plasma oxytocin levels across pregnancy and the postpartum period predict mother-infant bonding. Psychol. Sci. 18, 965–970 (2007).

32. Carter, C. S. & Keverne, E. B. The Neurobiology of Social Affiliation and Pair Bonding. Horm. Brain Behav. 299–337 (2002). doi:10.1016/B978-012532104-4/50006-8

33. Ainsworth, M. D. S. & Bell, S. M. Attachment, Exploration, and Separation: Illustrated by the Behavior of One-Year-Olds in a Strange Situation. Child Dev. 41, 49 (1970).

34. Boorman, R. J., Creedy, D. K., Fenwick, J. & Muurlink, O. Empathy in pregnant women and new mothers: a systematic literature review. J. Reprod. Infant Psychol. 1–20 (2018). doi:10.1080/02646838.2018.1525695

35. Aspell, J. E., Lenggenhager, B. & Blanke, O. Keeping in Touch with One’s Self: Multisensory Mechanisms of Self-Consciousness. PLoS One 4, e6488 (2009).

